# Indigenous ancestry and admixture in the Uruguayan population

**DOI:** 10.1101/2021.06.09.447750

**Authors:** Lucía Spangenberg, María Inés Fariello, Darío Arce, Gabriel Illanes, Gonzalo Greif, Jong-Yeon Shin, Seong-Keun Yoo, Jeong-Sun Seo, Carlos Robello, Changhoon Kim, John Novembre, Mónica Sans, Hugo Naya

**Affiliations:** Bioinformatics Unit, Institut Pasteur de Montevideo, Montevideo, Montevideo, 11400, Uruguay; Instituto de Matemática y Estadística Rafael Laguardia, Facultad de Ingeniería, Universidad de la República (UDELAR), Montevideo, Montevideo, 11300, Uruguay; Molecular Biology Unit, Montevideo, Montevideo, 11400, Uruguay; Centro de Matemáticas, Facultad de Ciencias, Universidad de la República, Montevideo, Montevideo, 11400, Uruguay; Precision Medicine Institute, Macrogen Inc., Gyeonggi-do 13605, Korea; Department of Human Genetics, Department of Ecology and Evolution, University of Chicago, Chicago, Illinois, 60637, USA; Departamento de Antropología Biológica, Facultad de Humanidades y Ciencias de la Educación, Universidad de la República (UDELAR), Montevideo, Montevideo, 11200, Uruguay; Agregado de Cooperación Lingüistica y Cultural de la Embajada de Francia en Uruguay, Montevideo, Montevideo, 11100, Uruguay; Bioinfomatics Institute, Macrogen Inc., Seoul, 08511, Korea; Precision Medicine Center, Seoul National University Bundang Hospital, Gyeonggi-do, 13605, Korea

**Keywords:** Human genomics, population genomics

## Abstract

The Amerindian group known as the Charrúas inhabited Uruguay at the timing of European colonial contact. Even though they were extinguished as an ethnic group as a result of a genocide, Charrúan heritage is part of the Uruguayan identity both culturally and genetically. While mitochondrial DNA studies have shown evidence of Amerindian ancestry in living Uruguayans, here we undertake whole-genome sequencing of 10 Uruguayan individuals with Charruan heritage. We detect chromosomal segments of Amerindian ancestry supporting the presence of indigenous genetic ancestry in living descendants. Specific haplotypes were found to be enriched in ‘Charrúas’ and rare in the rest of the Amerindian groups studied. Some of these we interpret as the result of positive selection, as we identified selection signatures and they were located mostly within genes related to the infectivity of specific viruses.

Historical records describe contacts of the Charrúas with other Amerindians, such as Guaraní, and patterns of genomic similarity observed here concur with genomic similarity between these groups. Less expected, we found a high genomic similarity of the Charrúas to Diaguita from Argentinian and Chile, which could be explained by geographically proximity.

Finally, by fitting admixture models of Amerindian and European ancestry for the Uruguayan population, we were able to estimate the timing of the first pulse of admixture between European and Uruguayan indigenous peoples in 1658 and the second migration pulse in 1683. Both dates roughly concurring with the Franciscan missions in 1662 and the foundation of the city of Colonia in 1680 by the Spanish.

## Introduction

At the time of European contact with the peoples of present-day Uruguay (estimated in 1516, date of the landing of first Spanish conqueror Juan Díaz de Solís), the land was populated by several different indigenous groups for which there is varying and scarce data (Arce 2015). Historians have progressively reduced the number of described ethnic groups from three or four. The main one, the macro-ethnic Charrúa group (expression first proposed by Vidart (Vidart 1973) to describe the former Pampeans), was composed by groups known as the Charrúa and Guenoa and other minor groups (eg. Bohan and Yaro), while the Guaraní, an indigenous group more abundant in regions farther north from Uruguay and present-day Paraguay, arrived at the territory probably few centuries before Spanish conquerors (Cabrera 1989).

Charrúas, even though they participated in the independence war against Spain, were continuously persecuted, including by the Uruguayan government after independence (1830). The ‘‘Salsipuedes’’ massacre, in 1831, has been recorded as the final episode of the persecution of the Charrúa by the Uruguayan army, where the government of the country deliberately killed the majority of them. Only few indigenous men were able to escape, while women and children were taken as prisoners and distributed as servants across the country. The genocide, or extermination, of this ethnic group, has been an object of different historic and ethnohistoric studies (Acosta y Lara 1961, Acosta y Lara 1985, Acosta y Lara 1989).

Originating further north, in present-day Paraguay, the migration of Guaraní to Uruguay has been considered the first migration wave into Uruguay in historical times (Gonzalez 1989). The last recorded arrival was in 1829 when General Fructuoso Rivera (first president of the Uruguayan Republic) brought several thousands of natives from the Jesuits Missions (probably mostly Guaraní) to found two cities, Santa Rosa del Cuareim and lastly, San Borja del Yi. In 1767 the Jesuits Missions were expelled from the territory, leaving thousands of individuals living in those regions (Curbelo 2009, Curbelo Barreto 2010).

Despite Guaraní’s presence, Uruguayan national identity has been related to the disappearance of the Charrúas and for many years, it was believed that it was a ‘‘native-free’’ country. After the genocide, the indigenous contribution was overwhelmingly underestimated: in the first National Census (1852) indigenous peoples were not even mentioned (categories were only “Whites”, “Mulattoes”, “Blacks”, “foreigners”); The most recent National Census (2011) introduced questions about ancestry and ethnicity, revealing that only 4.9% of the population self-report as having at least one indigenous ancestor, though contemporary studies on mitochondrial genomes have revealed indigenous haplogroups in frequencies of 31-37%, and the nuclear genome provides estimates of 10 to 14% indigenous ancestry (Sans et al 1997, Hidalgo et al 2005, Bonilla et al 2004). At present, Uruguay is experiencing a reemergence of the Charrúas cultural identity (Magalhaes Michelena 2017, Rodriguez Verdesio 2017), even though it is not necessarily linked to ancestry.

While admixture has been sufficient and not denied by any present association or self-recognized ethnic group, full Uruguayan indigenous genomes are lost. However, as shown by studies of other Amerindian ancestries in Latin America (e.g. Schroeder et al 2018) the reconstruction of indigenous gene pools using admixed gene pools opens a new tool to understand the origins of the ancient inhabitants of America. Moreover, they can be used to identify genomic segments that could belong to different ethnic groups, interesting to history and individual and national identity. As Bodner et al (Bodner 2012) states, data about indigenous or mixed South American populations will allow a more detailed panorama of migrations inside the continent. Admixed or urban populations have proven that they are useful predictors of indigenous diversity, at least considering mtDNA (Tavares 2019).

Studies of ancient DNA have also contributed to the understanding of past populations of the continent through the recovering of extinct genomes, as well as to prove continuity from past to present times. At present, several complete genomes from prehistoric indigenous groups have been sequenced, eg. the Pleistocenic rest from Anzick, Montana (Rasmussen et al 2014), the Ancient One/Kennewick Man (Rasmussen et al 2015), four individuals from the Northwestern coast (Lindo et al 2017), as well as 15 from different parts of America (Moreno-Mayar et al 2018). More recently, a database of over a thousand European ancient genomes was released (Mathieson et al 2018, Olalde et al 2018). In Uruguay, three ancient mitogenomes have been published. One of these belonged to the mtDNA haplogroup C1d3, which has been found only in Uruguayans and established continuity from an individual dated to 1610±46 years before present with one Charrúa Indian from the 18-19th centuries, and present population individuals (Sans et al 2015), while the other two lineages, belonging to haplogroup C1b, seem to have no continuity in present times (Figueiro et al 2016).

Moreover, the lack of prehistoric remains that can be identified with ethnic groups living in historical times, as well as the lack of historical remains which belong to indigenous groups, will be an obstacle to use them as the unique approximation to the knowledge of Amerindian genetic characteristics. Hence, the possibility of identifying indigenous segments of DNA from present population will be a useful proxy to reconstruct indigenous genomes and, when possible, to identify different indigenous ethnic origins.

In the present work we report whole genome sequences of 10 individuals with known (self-declared) Charrúas ancestry to gain insight to the indigenous ancestry of Uruguayans. Genome wide ancestry proportions, differences in self-declared and genomic estimation, similarity to other indigenous tribes in the region and selection signatures within indigenous tracts were determined. Additionally, the timing of early admixture with Europeans and Africans was estimated from the genomic data, and concurs with historical events.

## 2. Results

### 2.1 High indigenous signal is observed in Uruguayans with self-declared indigenous ancestry

Ten individuals with self-declared known Charrúas ancestry were included in the study. Family information and historical records were validated to include each sample. Samples had a very high quality as seen by the high throughput of each run (mean of 801542015 reads), low number of duplicated reads, high percentage of mapped reads (mean of 94.81% on the de-duplicated reads) and high percentage of reads with QC>30 (mean 83.95%).

Principal component analysis (PCA) of the samples was carried out using as a reference set individuals from the 1000 Genomes Project (The 1000 Genomes Project Consortium 2015), Simons Diversity Project (Swapan et al 2016).

In the PC1/PC2 space, Charrúas descendants were placed on the European-Asian/Amerindian axis (blue triangles), consistent with a high degree of admixture degree (figure 1A). In higher PCs (Figure S1), the ‘ Charrúas’ samples locate more closely towards other Amerindian reference samples than Asian samples. Hence, it is shown that the sampled individuals have admixture with Amerindian, proving that there is still indigenous signal in this population. Figure 1C shows global ancestry estimations using ADMIXTURE (K=4) of the 10 individuals, which are consistent with the PCA, showing a significant indigenous signal on the Uruguayan genomes (light blue color bars; higher K values are shown in Figure S2; comparisons with different methods are detailed in Supplementary material S3).

**Figure 1.**
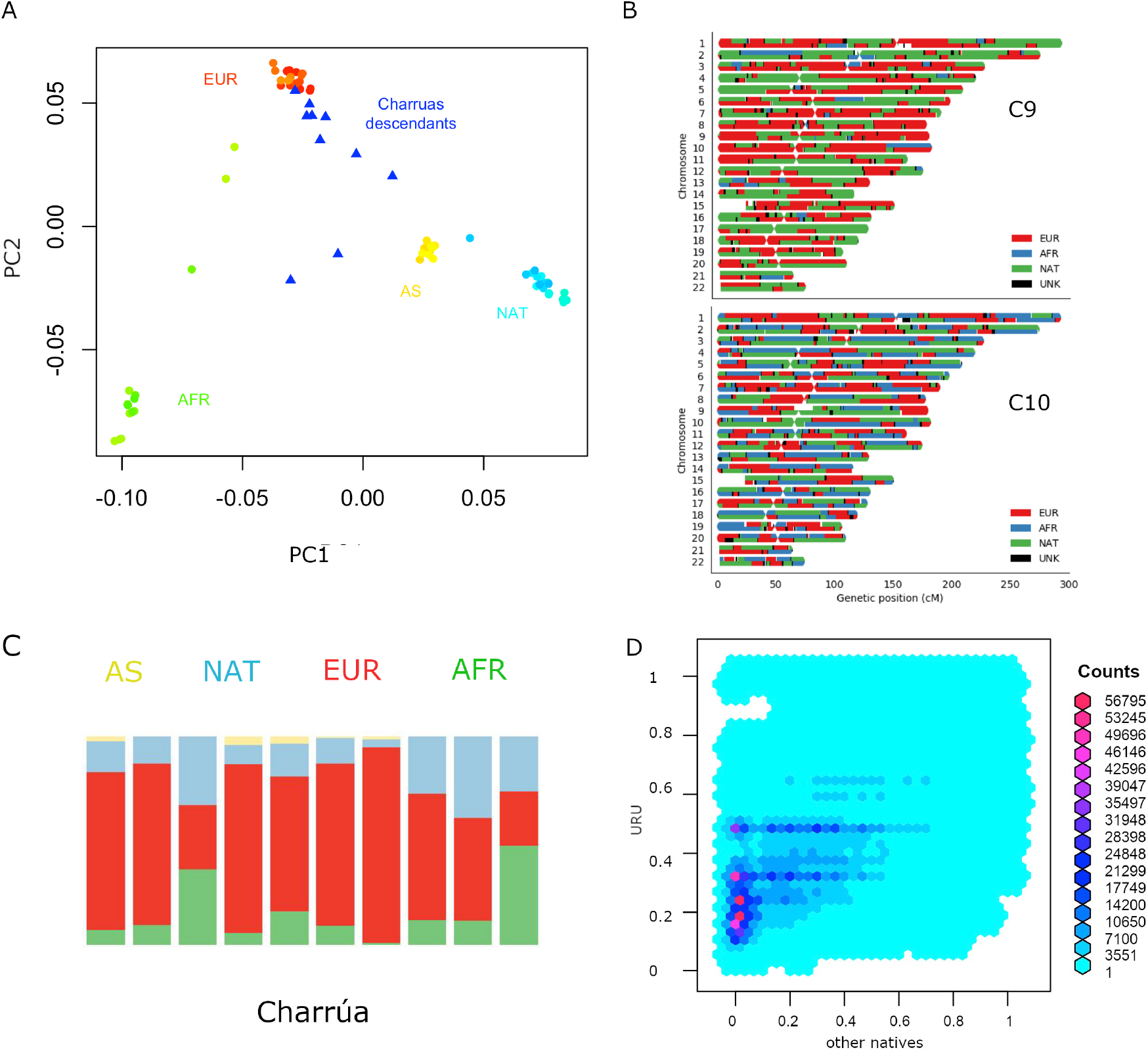
Admixture of Uruguayan samples. A. PC1 against PC2 of a PCA with complete genomes from different populations: Amerindian (cyan), Europeans (red), Asians (yellow), Africans (green) and ‘Charr\’ua’ descendants (blue triangles). B. Karyograms for two sample individuals showing results from local ancestry inference using a three-way model of admixture. Colors indicate the most probable local ancestry assignment (red: European, blue: African, green: Amerindian). C. Admixture plot corresponding to k=4. Reference populations were the same as in A: Africans (green), Asians (yellow), European (red) and Amerindian (light blue). 10 bars corresponding to the ‘Charrúas’ descendants. D. Frequency comparison between haplotypes coming from Uruguayan Amerindian and the rest of the indigenous groups. Reddish hexagons, correspond to a high amount of haplotypes corresponding to that pair of frequencies. Cyan and blue hexagons correspond to less haplotypes for that pair of frequencies.

We next carried out local ancestry inference using a three-way model of admixture using RfMix. Figure 1B shows local ancestry estimations for two example individuals (one with the most Amerindian ancestry and the other with the most African ancestry; full results, Figure S4). One of the individuals shows a considerably high proportion of indigenous ancestry (top, in green), while the other has a more even contribution of all three ancestries (bottom, AFR, EUR and NAT).

Additionally, we determined mitochondrial and Y-chromosomes haplogroups. Six of the mitochondrial haplogroups found are typical of living Amerindian individuals: B2e, B2 (n=2), B2b, C1d and C1b; three European: J1c1f, V, J1c and one African: L1b1a. None of the canonical Amerindian haplogroups for the Y chromosome were observed.

### 2.2 Frequent ‘Charrúas’ variants and haplotypes are rare in other populations

A total of 4524317 genome-wide variants (according to reference genome GRCh37) were found within ‘ Charrúas’ tracts (at least present in one individual). Of those variants (within indigenous tracts), 809497 were highly frequent in observed ‘Charrúas’ tracts (frequency>0.8), and of those, 105 were very rare in the rest of the world (frequency <0.01 in 1000Genomes and gnomAD). Table S5 compares the frequency of all ‘Charrúa’ variants with the rest of the world (1000G). The 105 globally rare variants fall into the following functional categories: 62 intergenic, 32 intronic, 6 ncRNA intronic, two downstream (variant overlaps 1-kb region downstream of transcription end site), one lncRNA, one 3’UTR and one 5’UTR.

Of the original 4524317 variants only 26628 were exonic and the rest non-coding, including splicing, UTRs, intronic, intergenic, etc. Of the total, only one was novel (not reported in dbSNP version 151) in chromosome 14 and genomic position 48780303, an intergenic variant. We also assessed patterns of haplotypic variation. Haplotypes were defined on the basis of nonoverlapping windows of 100 variants and their frequency were determined for the ‘Charrúa’ population and the rest of the indigenous groups.

Figure 1D compares the frequency of such haplotypes in ‘Charrúa’ versus all other indigenous groups. Enrichment of reddish hexagons are seen with low frequencies in other indigenous groups and in higher frequencies in ‘Charrúa’. This implies that ‘Charrúa’ have characteristic, private haplotypes, which are often not present in the other indigenous groups.

In order to see if some of these haplotypes may be the result of recent positive selection, we analyzed signatures of selective sweeps within the ‘Charrúa’ individuals. Taking only the indigenous tracts into account, iHS analysis was performed (Voight 2006).

Figure 2A shows the log transformed p-values of the iHS analysis for each “indigenous” position in the genome. The locations of the top 30 iHS values are reported in Table S6, and we further investigate the top two of these (marked with an asterisk in figure 2A).

**Figure 2.**
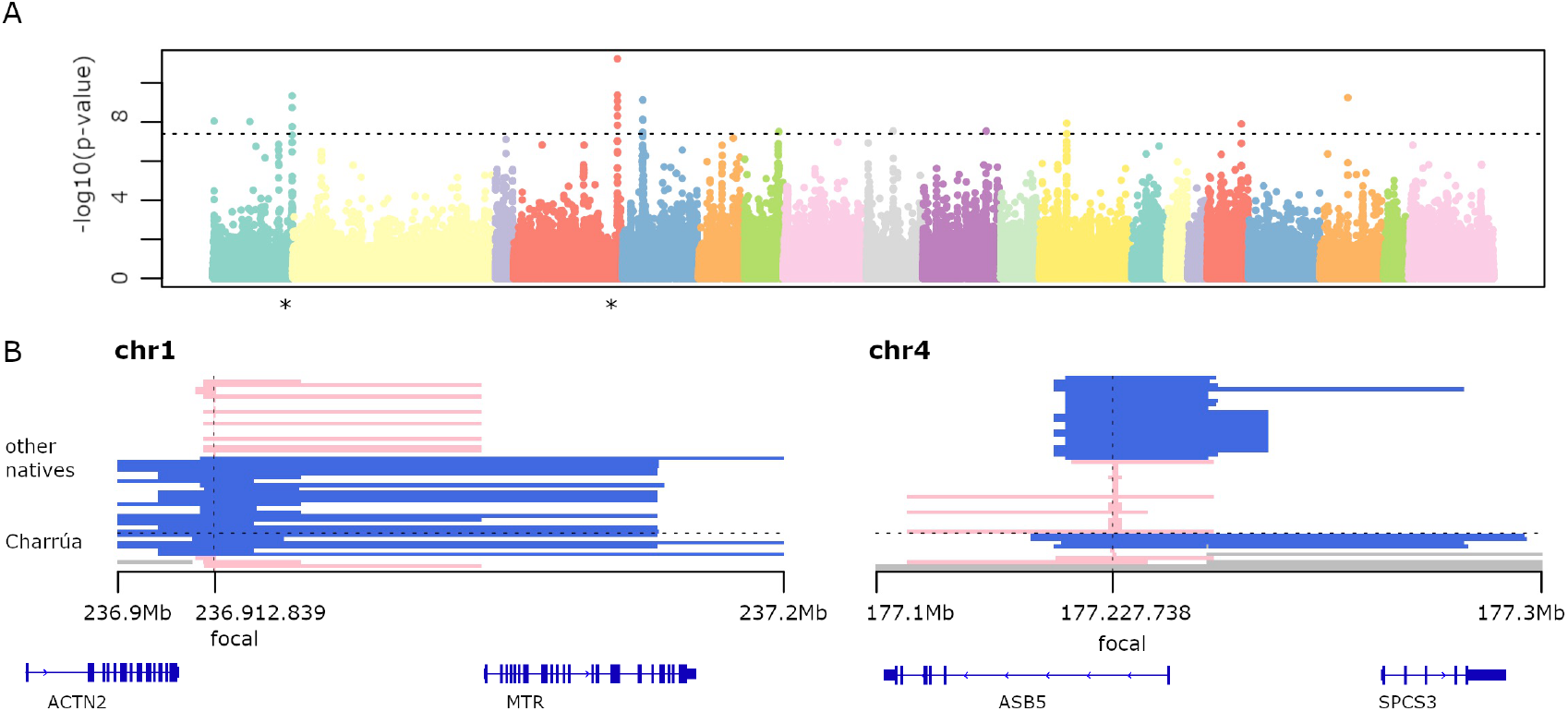
Selection signatures. A. −log10 of p-value of genome-wide iHS analysis. Dots above the line correspond to significant p-values (-log10(p-values) > 7). Several regions are found to be significant. Those marked with the asterisk are further analyzed. B. Significant haplotypes according to iHS. To the left, is the region in chromosome 1. Blue haplotypes are conserved among 6 out of 10 Uruguayan haplotypes, and its extension is shown with the bars (ranging from 236.9Mb to 237.2Mb). Four individuals have a different short haplotype (in pink). To the right, a highly conserved blue haplotype among Uruguayans (4 out of 9), which is broken in other Amerindians (shorter blue bars). Missing data is represented in gray, corresponding to masked haplotypes. Also, genes falling within those regions are shown (bottom).

One is found on chromosome 1 (0.3Mb long), which is present in 6 out of 10 Amerindian haplotypes. It is also found in the rest of the indigenous groups: Karitiana (4), Quechua (1), Zapotec (1), Mixe (3), Mayan (1), Piapoco (1), Chane (1).

Genes falling in that region are ACTN2 (only partially, the 3’ end) and the complete MTR gene. ACTN2 gene encodes a muscle-specific, alpha actinin isoform that is expressed in both skeletal and cardiac muscles. It is related to RET signaling and signaling by GPCR pathways (Rebhan 1997). The MTR gene encodes the 5-methyltetrahydrofolate-homocysteine methyltransferase. This enzyme catalyzes the final step in methionine biosynthesis and is involved also in folate metabolism (Rebhan 1997).

The second large iHS value region was found on chromosome 4 with a 0.2Mb extended haplotype (figure 2B, right panel), that is found in 4 out of 9 observed ‘Charrúa’ haplotypes (see section 5.4)

The extended haplotype is also found in Karitiana haplotypes (5), Quechua (2), Zapotec (3), Mixtec (1), Mixe (2), Mayan (3), Pima (1), Piapoco (2), Surui (2) and Chane (1).

The other 4 Uruguayan individuals have a smaller haplotype, and it is conserved among a smaller fraction of indigenous groups: Quechua (1), Maya (1), Chane 1). Genes falling within that region are ASB5 and SPCS3. ABS5 is a protein coding gene, related to the pathways of the Innate Immune System and Class I MHC mediated antigen processing and presentation. SPCS3 is also a protein coding gene, related to viral mRNA translation and gene expression (Rebhan 1997). To investigate whether the low coverage of indigenous tracts per SNP in the calculations of the iHS values might generate artificial extreme values, we compared the fraction of observed Amerindian haplotypes per locus with the calculated iHS values (figure S6).

Even though we observe a range compression in the highest frequencies (from 0.7 to 0.8), there is no general tendency of range compression from the lower coverage frequencies towards the intermediate frequencies. Hence, there is no obvious bias in the results due to the uneven/low Amerindian coverage. A similar approach was applied in (Gautier `and Vitalis` 2011), by using the proportion of a specific ancestry to help in the detection of selection signatures.

### 2.3 Correlation between self-declared ancestry and genomics estimations

The average self-declared indigenous ancestry in the sample was 15%. When compared to the genomic estimation of ancestry, the self-declared ancestry is on average lower (figure 3A, paired t-tests p-value<0.03).

**Figure 3.**
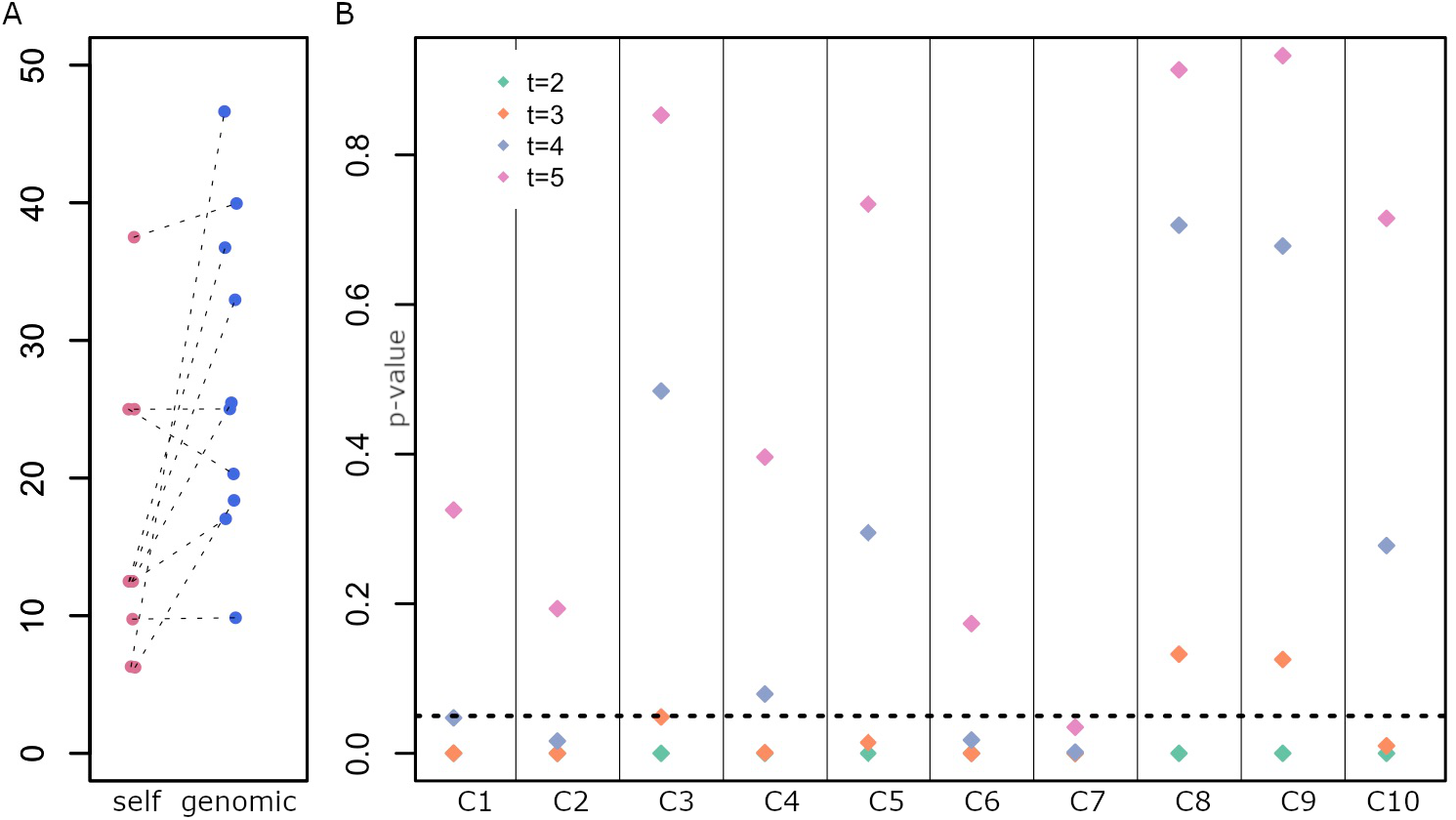
Determination of the generation of the last full indigenous ancestor. A. Distribution of self-declared Amerindian ancestry (self) and genomic estimates (genomic). Each dot is an individual. B. Test of a model with a full indigenous ancestor in each generation for each individual. P-values for rejecting the null hypothesis are on the y-axis, and individuals are displayed along the x-axis.

Five individuals have a relatively low self-declared Amerindian ancestry and a high genomic estimation, with a mean difference between both ancestry estimates of 22%. Those individuals potentially have indigenous ancestry from both matrilineal and patrilineal branches. Figure 1B shows the karyogram of two individuals, where homozygous tracts of indigenous ancestry along the whole genome can be observed (similar homozyous tracts are found for other individuals, Figure S2). Additionally, two individuals who declared European maternal ancestry have mitochondrial DNA from Amerindian origin.

Given the difficulty to identify the last ancestor with full indigenous ancestry for each individual from historical records, we developed a test (see 5.5) that tests, for each generation, whether the individual has a full indigenous ancestor at that generation or earlier in time. For each individual and each generation, we tested whether a full indigenous ancestor exists (null hypothesis). If p-values are smaller than 0.05, the null hypothesis is rejected, hence there is a low probability of the observed data if a full indigenous ancestor was present at that generation (for that individual).

Figure 3B shows the p-value for rejecting the null hypothesis for each generation (t=2, 3, 4 and 5) and each individual.

While for most individuals, we reject models where the full ancestor was 2 or 3 generations ago (C4, C5, C1), suggesting an ancestor around ~1830 or earlier, e.g., a great-great grandparent. In some cases, we could not reject an ancestor 3 generators ago (C3, C8, C9), consistent with a more recent ancestor (e.g., towards the end of 19^th^ century, a great grandparent). These results agree with the genomic estimations of Amerindian contributions (C3, C8, C9 have the highest indigenous ancestry, Figure 1C).

### 2.4 Inference of European contact and subsequent migrations

Since recombination events in each generation break down chunks of genome coming from the same parental chromosome, the length of resulting tracts assigned to distinct ancestries in admixed genomes might be informative of the migration modes and times (Pool Nielsen 2009). To explore the timing of European and African contact with the Amerindians in the region, the length distribution of continuous ancestry specific tracts of the ten individuals was considered. According to historical records, the Europeans (Spanish and Portuguese) came first to the present-day Uruguay, followed by probably more Europeans along with Africans, who were brought as slaves. Hence, in this study, two different migrations scenarios were evaluated: i) EUR,NAT + AFR, meaning a first contact of Europeans and Amerindians, followed by a pulse of Africans, and ii) EUR,NAT + AFR + EUR, meaning European and Amerindian contact, followed by a pulse of Africans, followed by a subsequent pulse of European migrants. Model ii) has two additional parameters, corresponding to time and proportion of the subsequent European migration.

Table 1 shows the log-likelihood for both fitted model, with estimated parameters including the number of generations ago for admixture (GA) for both models. We performed a likelihood ratio test to compare the nested models, 2 * (174.26-171.56) = 5.4, which under a chi-square distribution (df=2) has a p-value of 0.033. Hence, we selected the model ii) for further consideration (figure 4).

**Figure 4.**
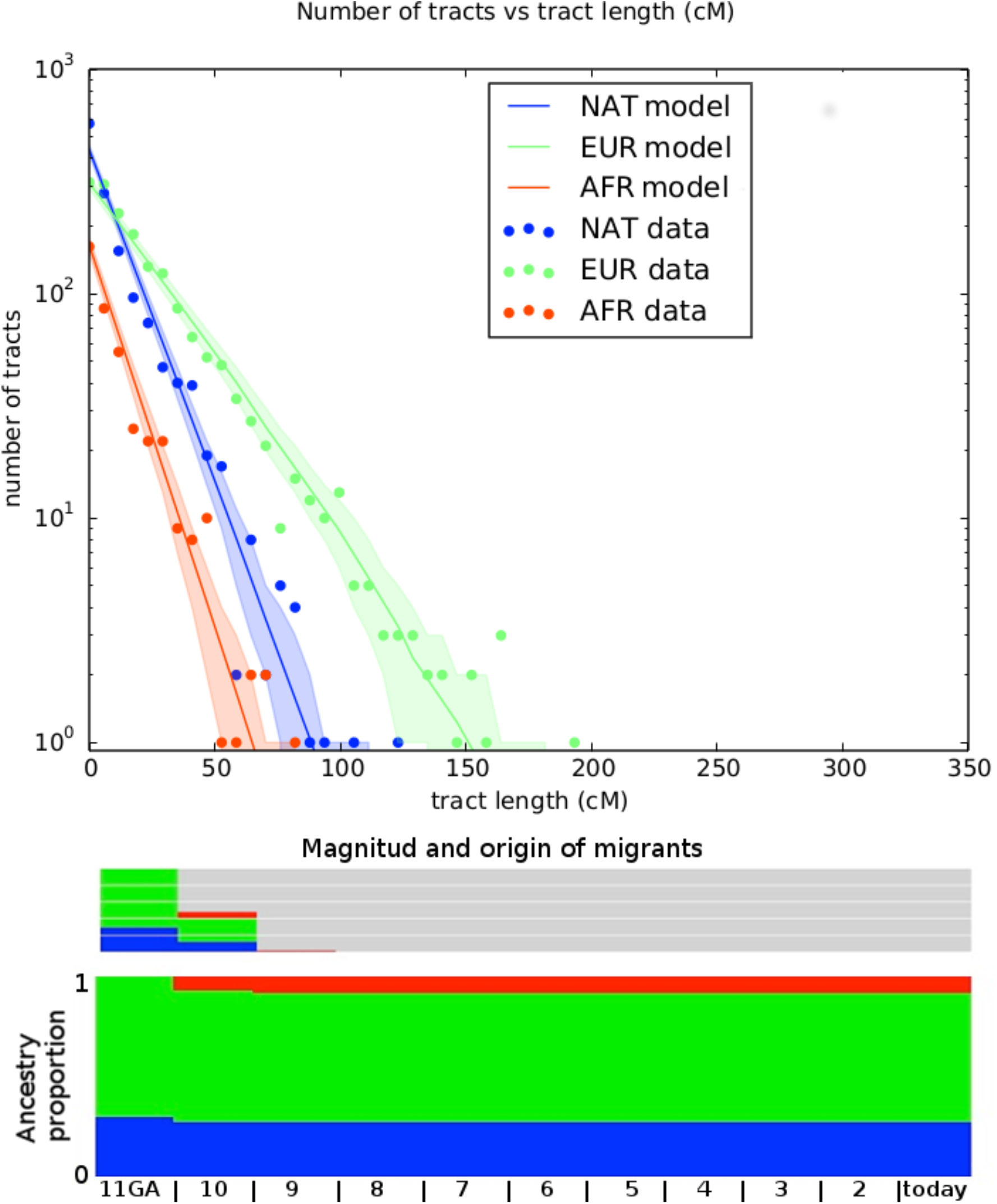
Representation of best fitting migration model. Top panel shows the number of tracts vs. the tract length. Observed distribution of ancestry tracts is represented by scatter points, while the solid-colored lines represent the distribution of the model, with shaded areas indicating 68.3\% confidence intervals. Models implemented in TRACTS were used to find the best model that fits the observed data. Admixture time estimates (in number of generations ago), migration events, volume of migrants, and ancestry proportions over time are given for each population under the best-fitting model.

**Table 1:** Two migration models were tested. i) EUR,NAT + AFR is a simple model were NAT has first contact with EUR in time T_1, with a subsequent pulse of AFR migration in T_2; ii) EUR,NAT + AFR + EUR is model with a first NAT and EUR contact at time T_1, followed by an AFR pulse at T_2 and with a subsequent EUR pulse at T_3. Log-likelihood of each model is given. TRACTS estimates the generations (in GA units) of the first NAT-EUR contact, which is given under GA.

|  | EUR,NAT+AFR | EUR,NAT+AFR+EUR |
| --- | --- | --- |
| log-likelihood | -171.56 | -174.26 |
| αNAT_prop | 0.29 | 0.29 |
| τtime_(EUR, NAT) | 8.3 | 8.5 |
| τtime_(AFR) | 6.7 | 7.8 |
| τtime_(EUR2) | NA | 0.36 |
| GA | 11 | 11 |

The migration matrix is represented in the bottom panel of figure 4 and reveals that the first European contact with the Uruguayan ‘Charrúas’ was 11 generations ago, which corresponds to around the mid-end of XVII century (1658 with the formula 1958 – (GA + 1) * 25 as in (Fortes-Lima 2017), being 1958 in this case the median of all participants’ birth dates. GA: generations ago). The second European migration contributed 30% to the population and was 10GA ago (1683). The first African pulse was 10 GA (1683), with an ancestry contribution of 7%. The second one was 9 GA with an ancestry contribution of 1%.

To further investigate the contribution of different migrants to the Uruguayan Amerindians we determined the expected heterozygosity according to the ancestral contributions. For this we did a linear model with the ancestral proportions as the independent variables and heterozygosity as the observed variable (see section 5.7). The calculated coefficients for the Amerindian, European and African, are 0.29, 0.31 and 0.46, respectively (Figure S7B, first left panel). Hence, the African ancestry component has the larger effect on the heterozygosity of the Uruguayan Amerindians individuals studied here. Also, we compared the heterozygosity of individuals with full European, Amerindian and African ancestry, with the determined heterozygosities by the linear model (for each ancestry component) of Uruguayan indigenous descendants. We observed that those European and Amerindian heterozygosities in Uruguayans are comparable with those of individuals of that ancestries. Sardinians have almost the exact same heterozygosity as the European component of Uruguayans and, the “Amerindian” heterozygosity of Uruguayans lies between Mayan and Karitiana. However, African heterozygosity in Uruguayan individuals is larger than African individuals such as the YRI, LWK samples (Figure S7A). This means, on the one hand, that the high heterozygosity in Uruguayan individuals is mostly due to the African ancestry and on the other, that African component in Uruguayan sample in highly diverse.

### 2.5 Genomic similarity between ‘Charrúa’, Kaingang, Diaguita and Guaraní

The similarity of ‘Charrúas’ to other South American tribes and the history of admixture for ‘Charrúas’ are unknown. The only available information comes from relatively scarce historical records and oral histories. To assess similarity between ‘Charrúas’ and other South American tribes we used the data collected here along with that of (Reich et al 2012). The latter is a merged dataset describing 525 individuals from different populations (214 European, 109 African, 202 Amerindian), including 169 individuals from 14 South American populations, with a final genotype set covering 364K SNPs. Guaraní (from the Paraguay-Brazil and Paraguay-Argentina border), Aymara, Quechua, Chilote, Diaguita, Kaingang, Wichi, Toba, Guahibo, Piapoco, Karitiana and Surui were some of the represented populations.

To analyze the ancestry components in the ‘Charrúa’ descendant genomes collected here, the ADMIXTURE method was used, with K ranging from 3 to 19. The higher the number of clusters K, the finer the structure that can be analyzed. With K=19 all ‘Charrúas’ had a high contribution of three mostly European clusters (predominately represented by TSI, Sardinian and Russian individuals) and two mostly African clusters (predominantly, YRI and LWK, Figure S2). The 14 remaining clusters correspond mostly to indigenous groups. Figure 5A shows the classical admixture plot (with K=19) but without the European/African contribution, to allow a closer inspection of the Amerindian ancestry of our sample. A shared cluster is observed in almost each ‘Charrúa’ individual (indicated in blue in the figure). That cluster is mainly shared by Wichi, Ticuna, Toba, Guahibo and additional groups with more complex ancestry (Piapoco, Chane, Guaraní, Diaguita and Yaghan). A cluster (light orange) shared with Andean groups (Aymara and Quechua), is present in some ‘Charrúas’ descendants but in a smaller proportion than the blue one. Only three Charrua descendants have a non-negligible contribution of a cluster (green), which is shared mostly with Mexican indigenous individuals (Zapotec, Mixe, Mixetec, and Maya) though present also in small amounts in indigenous groups geographically closer to Uruguay (e.g., Diaguita). Several of the individuals share some ancestry with two clusters (dark orange and peach), that is shared with Amazonian groups (Karitiana and Surui). Finally, a small proportion of a cluster (purple) shared with Pima individuals is found in several of the Charrúa descendant individuals. These results show that this sample is generally homogeneous with respect to its indigenous genetic background (One individual, C7, is a visual outlier, though we note the ancestry proportions may be more difficult to infer as this individual was inferred to have ~9% indigenous ancestry).

**Figure 5.**
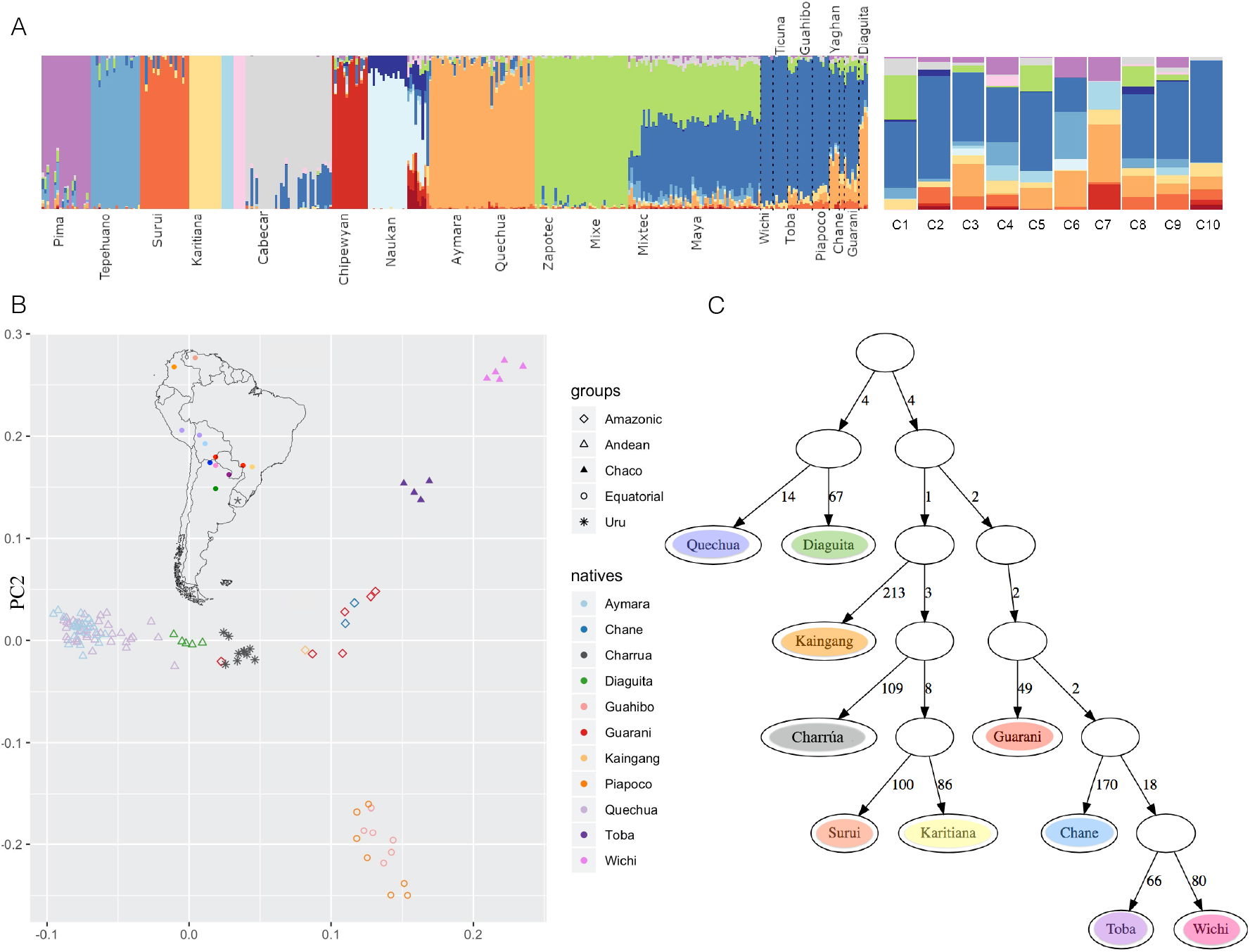
Comparison with other indigenous groups. A. Admixture at fine scales (with K=19). After running ADMIXTURE for K=19, the European and African clusters were discarded, and only indigenous contributions were considered. To the left, the reference Amerindian panel is shown and to the right the 10 Uruguayan indigenous descendants, where a high proportion of blue is observed in most of them. B. PCA admix on masked chip data C. qpgraph on masked chip data. The number on each edge represent genetic distance units scaled by a factor of 1000.

Additionally, we did a PCA on the Amerindian ancestry tracts using PCAadmix (Brisbin et al 2013) with local ancestry called by RFMix (figure 5B). The PCA shows ‘Charrúa’ descendants located between the distribution of Guaraní/Kaingang (red/light orange diamonds) and Diaguita (green triangles) individuals, suggesting closest similarity with those groups. Wichi/Toba and Guahibo/Piapoco groups remain in distant positions in the PCA space with respect to the ‘Charrúa’ descendant individuals. Moreover, the group Kaingang, Guaraní and Diaguita had the lowest F_ST_ values with the ‘Charrúa’ descendants (Figure S8).

Using the same data, admixture graphs were estimated with qpgraph (Patterson et al 2012). Figure 5C shows a plausible graph for the current data set (Z score = −3.197, corresponding to the following D-statistic outlier: [Surui, Kaingang], [Wichi, Guaraní]). Admixture branches did not improve the score. In the graph the ‘Charrúa’ are positioned among current Brazilian Amerindian groups; specifically, the ‘Charrúa’ are an outgroup to a subtree with Karitiana and Surui, with Kaingang as the most immediate outgroup to the Charrúa, Surui, Karitiana clade. The branch lengths (measured in units of genetic drift) suggest a star-like topology where many of the ancestral nodes are close to the root.

## 3. Discussion

We found a substantial indigenous genomic contribution in the ‘Charrúas’ descendants that participated in this study, despite there are no isolated indigenous communities living in Uruguay in the present.

These individuals have twice the estimated indigenous ancestry as the expected for the general Uruguayan population and is in most cases also higher than the estimations for Montevideo (around 15%), where the individuals were sampled (Hidalgo et al 2005, Bonilla 2015) (ancestry estimations range from 9\% to over 40\% with a mean of 27\%).

In general, the overall level of Amerindian ancestry was higher than expected from self-report in this sample. Particularly, in most cases, individual genomic data was consistent with indigenous ancestry on both parental lines, which was remarkable as each participant self-reported indigenous ancestry only from a single paternal side. This is consistent with the underestimation of indigenous ancestry in the general Uruguayan population (Sans et al 2011), and is perhaps notable that this underestimation occurs also in this set of individuals that self-identify with Charrúan heritage. Two individuals also had a high proportion of African ancestry (0.21 and 0.30) though African ancestry was generally not reported by the participants, probably meaning that their sense of identity was more related to their Charrúan ancestry, or the admixture event was sufficiently in the past to be unknown to them.

As previously shown, in the post-colonial period matings that occurred in Latin America were mostly between Amerindian women and European men (Ongaro et al 2020, Sans et al 2000). In addition, indigenous genocide in Uruguay (‘‘Salsipuedes’’) exterminated the majority of ‘Charrúa’ men. Thus, we found indigenous mtDNA haplotypes in 6 out of the 10 individuals, while none of the five men had an indigenous Y-chromosome.

The indigenous mitochondrial haplogroups found in this sample were B and C, which are typically found in Uruguay (Bonilla et al 2004, Bonilla et al 2015}). Figueiro and Sans (Sans et al 2015) discussed the difference of origins between the haplogroups A, B and C, with A being more commonly found in Amazonian peoples (particularly Guaraní), and B and C more associated with Uruguayan prehistoric sites and possibly Argentinean Chaco or Pampa (in spite of having a broad distribution).

A scan for recent positive selection (using iHS) showed several regions that might be under selection.

A selection signal was observed in chromosome 4 spanning the ASB5 and the SPCS3 genes. The SPCS3 (Signal peptidase complex subunit 3) gene plays an important role in virion production of flaviviruses such as West Nile virus, Japanese enchephalitis virus, Yellow Fever virus, Zika and Dengue virus type 2 (Zhang et al 2016). The latter two, most likely present in the Uruguayan region before colonization. A subset of endoplasmic reticulum-associated signal peptidase complex proteins was necessary for proper cleavage of the flavivirus structural proteins (prM and E) and secretion of viral particles. Hence, those proteins (among them SPCS3) are required for flavivirus infectivity, and these were associated with endoplasmic reticulum functions including translocation, protein degradation, and N-linked glycosylation. The gene region within the selected haplotype, which is present in 4 out of 9 Uruguayan haplotypes, presents almost no variants, meaning that mostly the ancestral allele is conserved. Only one variant is found at position 177253006, which has no frequency reported in the gnomAD database (Karczewski et al 2020}. If this variant could have implications in immune system evolution and adaptation for Uruguayan indigenous groups cannot be stated without further studies.

A region in chromosome 1 was also found, that is highly frequent in Uruguayan Amerindians and other indigenous groups. The genes included in this region are ACTN2 and MTR. The MTR gene is an enzyme that is involved in methionine biosynthesis. Methionine synthase helps convert the amino acid homocysteine to methionine (Oohora et al 2016, Liu et al 2010). The ACTN2 gene, is a muscle-specific, alpha actinin isoform that is expressed in both skeletal and cardiac muscles.

According to our estimated time frame of admixture, the first European-Amerindian was around 1660 and the second around 1680. In 1516 the Spanish Juan Díaz de Solís, reached the (now) Uruguayan coast for the first time. During the 16^th^ century, forts and villages were founded by the Spanish, however, none of them survived more than five years. Thus, no admixture persisting to today was expected during that period and this was confirmed by the genomic data. Interestingly, the first estimated admixture event concurs with the establishment of the Franciscan mission, Santo Domingo de Soriano, founded in 1662-1664. According to historical records, allegedly mostly Chaná (a different indigenous group with uncertain relationship to the Charrúas) lived in the Santo Domingo de Soriano Franciscan mission (Acosta y Lara 1961, Barreto et al 2011). This could imply that, either ‘Charrúa’ were also living together with Chaná and Franciscan individuals, or simply that we cannot distinguish between these indigenous subpopulations with our samples yet, or even that we cannot determine the exact time frame of the contact (a change in one generation and/or using a different assumed generation length can move the estimated date by more than two decades). In the same line, the second migration pulse in ~1680, tentatively concurs with the foundation of the Nova Colônia do Santíssimo Sacramento, that brought together according to historical records, more than 400 soldiers, around 60 African slaves, and some Amerindians (Azarola 1940).

Using the TRACTS model, the European genomic contribution in the first generation of admixture was estimated as 70% with 30% of indigenous contribution. This over-simplifies the probably real historical process, where Amerindian were the vast majority in the region (at least until 17^th^ and probably also 18^th^ century, but for the latter there is scarce data regarding population numbers), only few admixed with Europeans and the rest of the unadmixed Amerindian probably left no footprint in the genome of living descendants, most likely due to the massive genocide (Salsipuedes). Further, the TRACTS model fit a second European wave 10 generations ago (approximately 1683) that contributed 30% to the population already living in the territory. Additionally, the first inferred African pulse was also ten generations ago. These data may suggest that the first pulse of African slaves came here together with the Iberian soldiers. Even though historical reports show several thousand of slaves came after the foundation of Colonia, their genomic contribution is inferred genetically to be low, around 7%. This could reflex the social behavior during slavery times. A second African pulse was also estimated in 1708, contributing only 1% to the gene pool of the historical population. This may correspond to a further pulse of African slaves coming from further Spanish ships. In 1726 was the foundation of the city of Montevideo and even though, according to historical records, only Spanish participated in this process, a couple of years later due to the high activity of a port city, slaves were brought as workforce to keep up with the labor that the city required (Bracco et al 2012).

Uruguayan indigenous group is placed between Andean and Amazonic natives in the PCA. The closest populations from those groups are Diaguita (Andean) and Guaraní and Kaingang (Amazonic). This genomic similarity between these groups can be explained based on admixture events or shared ancestors. In particular, a Guaraní signal was expected, since different sources have shown that Guaraní arrived in Uruguay around the 13 ^th^ century, and they continue coming during historical times. Most of them came from the Jesuitic Missions, as mentioned before. Nevertheless, the signal was lower than expected, probably due to the heterogeneity (two different locations) and the small size of the Guaraní sample. Both facts could weaken admixture signals.

Kaingang, despite belonging to different linguistic families and having some degree of genetic differences (Petzl et al 1993), lived close in southern Brazil, a region also inhabited by Charrúa. Also, kidnaps and exchanges, especially of women, were common practices in different Amerindian groups during the historical period (Bracco 1998). These facts, together with close geographic distances, probably favored admixture events between Guaraní, Kaingang and ‘Charrúa’.

In contrast, the Diaguita similarity with Uruguayan indigenous descendants (and also to Guaraní and Kaingang), was at first, unexpected. However, the admixture between ‘Charrúa’ and Diaguita, especially from the Argentinean Northwest and Chilean “Norte Chico” (groups from the Chaco) is supported by several ethnohistorical records: Based on linguistics, Rona et al (Rona et al 1964) pointed out that the ‘Charrúa’ languages (as well as Minuán and Chaná) are related to Chaquean languages, specially the Vilela language group.

Additionally, Alcides D’Orbigny already in the 19th century, associated Charrúa together with Tobas and Mbocobíes for their customs, features and language (Acides 1839).

More recently, we found several records that support the Charrúa presence in the Chaco (Sanchez 2007). Between the Chaco and today’s Uruguayan territory, the “Argentinian Mesopotamia”, there is evidence of Charrúa presence in the 18^th^ century in the Cayastá’s reduction (Bracco 2016). Some studies even indicate that the expression “Band of the Charrúas” was used to describe the indigenous groups of the Paraná River (in Argentina) (Bracco 2004) and also, since the end of the 20^th^ century descendants of Charrúa have been establishing in the province of Entre Ríos (Argentina). Moreover, inter-ethnic admixture in the Jesuit reductions has been repeatedly described (Levinton 2009, Boccara 1998, Wilde 2009) and it could have nucleated different ethnic groups with an extensive geographical dispersion.

The admixture graph supports the similarity between ‘Charrúa’ and Kaingang, nevertheless Guaraní and Diaguita appear in different branches of the graph. However, those branch lengths (that are proportional to genetic drift) are small, meaning that a star-like phylogenetic structure could still be possible (only binary topologies are allowed in this approach). Admixture branches did not improve the score of the tree. Also, Diaguita-Guarani and Diaguita-Charrua have low F_ST_ values (Figure S8), which could also support early day differentiation with subsequent high amount of admixture events, or a later and recent differentiation of these populations. Additionally, Treemix results (Figure S9) support the relation between Charrúa and the ancestors of Diaguita (Andean group).

Not all tribes in South America are currently sampled. There is an under sampling of groups belonging to adjacent regions from the Charrúas, such as Chaná, Minuán and other groups. Hence, similarity and relations to those groups cannot be assessed. Additionally, the groups that were sampled are sometimes small (Kaingang, Chané) and sometimes sampled from very different locations, which can affect heterogeneity of the group, eg. Guaraní. In addition, a caveat is our small sample sizes of Charruan ancestry individuals and the amount of Amerindian information per individual.

The African ancestry component has the strongest contribution to the expected heterozygosity, and higher than expected for contemporary west African samples. Even though a high heterozygosity is expected in African ancestries, due to the evolution of populations within Africa, the calculated heterozygosity in Uruguayan sample is higher than African populations (Figure S8A). This could represent the diverse origin of African slaves that came to the territory. Conversely, the founders of European ancestry were most likely less diverse, since Spanish/Portuguese were the populations that invaded the territory. Even though African ancestry is in general low in this sample (0.08) some individuals have high contributions (0.30 and 0.21). For those, the more African ancestry, the highest the heterozygosity (Figure S8B, right and left panels). No correlation was observed for the other ancestries.

Regarding technical aspects, the relatively small indigenous signal of the genomes causes long stretches of missing data, where European and African ancestry were masked out. Also, this causes a frequency bias, since positions that are covered by few individuals have extreme frequencies, but this is due to sampling and not as a result of evolution. Either this can cause a bias in results (such as F_ST_) or a loss of information due to different strategies, such as filtering (only work with haplotypes that have at least a ‘reasonable’ number of haplotypes) or construction of pseudo-haploids, which were both used here.

## 5. Methods

### 5.1 Sample selection

Participants were selected after an interview with a social anthropologist. Interviews allowed to determine a high percentage of (expected) Amerindian ancestry and that participants were not related to each other. In general, participants declared to have at least a great grandparent or great great grandparent of Amerindian ancestry. Genealogies were constructed and several metadata information were gathered (ie. birth and death dates and places, occupation/craft, etc.) for these individuals.

15 participants enrolled in the study and 10 were randomly selected for whole genome sequencing, so that anonymity could be assured. All 15 participants (with or without WGS) were given the results of ancestry proportion inference (based on a custom SNP array of ancestry information markers, (Galanter et al 2012) as feedback for having participated in the study. Of the 10 selected for sequencing, six individuals declared to have ‘Charrúa’ ancestry, two declared a mixed (Guaraní and Charrúa) ancestry, one declared Guenoa and one unknown indigenous group. This information was blinded from the genetic analysis to assist in protecting anonymity of the participants. These procedures were approved by Institut Pasteur de Montevideo ethics committee with reference number IP011-17/CEI/LC/MB and with informed consent signed by each participant.

### 5.2 Sequencing and post processing of data

Ten whole genomes were sequenced in an Illumina HiSeq X ten with 30X coverage. Data was mapped with BWA (Li et al 2009) with default parameters, variant calling was performed according to the best practices with GATK (Mckenna et al 2010) and annotation with ANNOVAR (Wang et al 2010). A total of 10776269 high quality variants (SNPs and INDELS) were found among the 10 individuals.

Data was phased using Shape-it (Delaneau et al 2013).

MDS (using plink (Purcell et al 2007)) was performed on a merged data set of the Uruguayans, 1000G version 3 (The 1000 Genomes Project Consortium 2015) and Simons Foundations Amerindian individuals (Swapan et al 2016), resulting in 261376 variants after LD-pruning (using parameter --indep-pairwise 10 5 0.1).

### 5.3 Global and local ancestry estimations

Global ancestry estimations were done with ADMIXTURE \cite{alexander2009}, using as reference panel 1000G populations and 10 native genomes obtained from the Simons’ Foundation Project (Swapan et al 2016}. We used the same thinned data set as for the MDS analysis. In addition, in order to compare Uruguayan data set to a larger set of different indigenous groups, we used the SNP microarray data of (Reich et al 2012). In this study 52 Native Americans were genotyped at 364,470 single nucleotide polymorphisms, together with European and Africans, among others. The Uruguayan data set was assessed at those positions and was integrated to this data set. ADMIXTURE was run using this combined data set, using as reference unadmixed individuals as done in (Reich et al 2012).\\

In both cases (thinned data set with 1000G and chip data set) ADMIXTURE was run with K ranging from 3 to 18. For each K, an ad hoc assignment of the clusters labels was done. For K=3, the labels are most likely “European”, “African” and “Native”. This was used to calculate global ancestry proportions, in order to compare with RFMix results. For higher K’s the assignment of the labels is less straight forward, but can be made by analyzing the individuals present in each group.

Local ancestry estimation was done with RFMix (Maples et al 2013), using a reference panel that included 21 whole genomes of natives (Simons’ Foundation Project), 20 Africans and 23 Europeans from the 1000 Genomes Project (other set of parameters were evaluated, see Supplementary data S3).

### 5.4 Detecting signatures of selection

On the whole genomes of masked Uruguayan natives, we applied iHS calculations (Gautier et al 2012, Gautier et al 2017) for detecting intra population selection signatures. Each individual is considered as two independent haplotypes and we considered for the calculations only those haplotypes that had more than 15% of SNPs genotyped (min_perc_geno.hap=15) and SNPs genotyped on more than 40% percent of the haplotypes were considered (min_perc_geno. snp=40).

With these settings we were able to calculate an iHS statistic for 728935 sites. Considering an adjusted p-value of 4.12e^-8^, which corresponds to a family wise error of 0.03 with a Bonferroni correction 0.03/728935. The −log value is −7.4 (significance line in the figure 2A).

On both most significant peaks (chromosome 1 and 4) a visual inspection of the haplotype was made. Starting from the focal SNPs that iHS identifies, the haplotype is defined to the right and left. When one base differs from the extending haplotype, the extension stops and the haplotype breaks. After the definition of the haplotype, its presence in other natives was evaluated. Conserved haplotypes can be seen as long horizontal bars including many individuals (eg. blue haplotype in figure 2B, left) and haplotypes that are conserved in a population are seen as long bars in a subpopulation (eg. blue haplotypes at the bottom of figure 2B, right), and in other populations they appear as small centered bars.

The different colors in the figure correspond to different haplotypes that were extended from the focal SNPs.

### 5.5 Determining the generation at which an individual has a complete native ancestor with high probability

The objective of this hypothesis test is, fixed a number T of generations, and given the length of the maximum Charrúa haplotype for every non-sexual chromosome, assess if it is possible that one of the individual’s great^{T-2}^ grandparent is a complete Charrúa ancestor.

The test considers the family tree of the individual up to T generations in the past, regards the chromatids as intervals (measured in Morgans) rather than sequences of base pairs, and assumes the following meiosis model to generate a daughter chromatid from two chromatids for every pairing of the family tree:

i. Simulate the recombination points using a Poisson process with parameter L_i_, (length of the chromosome in Morgans). After adding the borders of the interval [0,L_i_], we obtain {xo = 0, x_1_, …, x_n_, x_n+1_=L_i_,}. As long as we measure the intervals in Morgans, using a Poisson process to simulate the recombination points is not a strong assumption for the model.
ii. Select a chromatid at random, and consider the segment tr_1_=[0,x_1_] in the selected chromatid. This will be the first segment of the daughter chromatid.
iii. Switch to the other chromatid, and concatenate the segment tr_2=[x_1_,x_2_].
iv. Iterate the last step until the length of the daughter chromatid is L_i_.

This is a model which is closer to the biological model than other widely used models, such as the Wright-Fisher model. However, this model does not fulfill the Markov property, so we need to simulate all the pairings of the family tree if we want to obtain an approximation of the distribution function of the statistic under the null hypothesis. This can only be achieved if we restrict the null hypothesis to a simpler scenario:

H_0_: a^0^ has exactly one pure Charrúa ancestor, T generations ago. The other ancestors are pure from some other ancestral population.
H_1_: a^0^ has no pure Charrúa ancestors, T generations ago.

If we consider a smart choice for the test statistic, H_0_ will become a borderline case of H_0_, meaning that the statistic will be stochastically smaller under H_0_. The test statistic requires a scaled score for every chromosome, and a way to combine them in a final test statistic. In this work, we choose to use the (scaled) length of the maximum tract for every chromosome, and consider the maximum of the scores as the test statistic.

More details and choices for a test statistic can be found in Illanes et al (manuscript submitted).

### 5.6 Migration models

The results of local ancestry estimations with RFMix (using as reference panel whole native genomes, and 1000G Africans and Europeans) were used as input for the TRACTS tool (Gravel et al 2012). Here, the length distribution of African, European, and Native American ancestry tracts along each chromosome is determined and an extended Markov model is applied to compare the observed data with predictions from different demographic models considering various migration scenarios.

As stated, several different models were tested, using predefined models provided by TRACTS and examples in https://github.com/armartin/ancestry_pipeline.

We tested three models: i) EUR,NAT + AFR, ii) EUR,NAT + EUR and, iii) EUR,NAT + AFR + EUR. For each, log likelihood was obtained, and bootstrapping was performed as follows: first 100 iterations were done, and if results was promising, further 100 were run, and so on until a maximum of 400. Model i) was run with 200 iterations, ii) with 100 and iii) with 400.

### 5.7 Analysis of Heterozygosity

Most of the individuals had the same amount of heterozygosity (around 0.355) with the exception of two individuals. To understand from which population it comes from, a linear model with heterozygosities as the dependent variable and the ancestry proportion (EUR, NAT, AFR) as the covariates was fitted.

HET = a*EUR + b*NAT + c*AFR

With the result of the linear model it is possible to write each individual’s heterozygosity as a linear combination of the expected heterozygosities of each ancestral population. We then compared the heterozygosity of each ancestral population in the Uruguayan one, with the heterozygosities of non-admixed populations, with different ancestries.

### 5.8 F_st_, PCAadmix and Adxmiture graphs

Uruguayan native genomes were assessed at chip positions. RFMix was run on chip positions including 525 individuals + 10 ‘Charrúa’, considering the Europeans, Africans and Natives from the chip as reference population. Native tracts were considered for these further analyses. Fst was calculated only for South American tribes: Toba, Guaraní, Kaingang, Wichi, Chane, Diaguita, Surui, Karitiana, Guahibo, Piapoco, Arara, Quechua, Aymara, Chilote and URU. Only variants with high coverage of native tracts were considered, since Fst calculations are sensitive to extreme frequency values. We tried different coverage values, and decided to have at least 6 native haplotypes. Four haplotypes gave similar results regarding Fst/f3 statistics but produced worse p-values (figure S12). A heatmap was used to visualized the results (Figure S10). For PCAadmix and admixture graphs (qpgraph), both from admixtools, we generated pseudo-haploids. For the positions in the genome with half-missing ancestry calls (ie. one haplotype native, the other one EUR/AFR, hence most of them) one retrieves the native allele and at positions where both alleles are natives, one randomly samples one of the two.

Those pseudo-haploids are used to generate the PCA to compare only native tracts of Uruguayans together with other natives from the chip.

The admixture graph was generated as follows. First a subset of 9 Southern Natives were considered to build a reasonable topology of the graph. Those topologies had 2 outliers with the same Z-score of 3. Next, we tried to add the ‘Charrúa’ branch, first without admixture and then with. The best fitting topology is the one presented in figure 5C. An equivalent one, with the same Z-score is in Figure S12.

## 6. Data access

Genomic data generated in this study is available at http://urugenomes.org/lovd/variants.

## 7. Acknowledgments

First of all, we want to genuinely thank the 15 indigenous descendants that generously donated their blood samples to participate in this study.

Also, we want to thank Mallick Swapan from Simons Genome Diversity Project for giving us access to the data and assisting us in the process.

We want to acknowledge Moria Sotelo for helpful discussions regarding the history of Charrúas and the interaction with other groups from an archeological perspective.

We thank Mariana Brandes for helpful scripting and figure design. and also Garrett Hellenthal for very helpful discussions.

This study was funded by BID (Banco Iberomericano de desarrollo) in the context of the URUGENOMES Project (Proyecto ATN / KK-L4584-JR ‘‘Fortalecimiento de las capacidades técnicas y humanas para las exportaciones de servicios genómicos’’). Additionally, partial support was obtained from the Agencia Nacional de Investigación e Innovación (ANII, Uruguay) FSDA_1_2017_1_143647 and FOCEM (MERCOSUR Structural Convergence Fund).

